# Adaptation to a novel predator in *Drosophila melanogaster*: How well are we able to predict evolutionary responses?

**DOI:** 10.1101/005322

**Authors:** Michael DeNieu, William Pitchers, Ian Dworkin

**Author notes:** Data and Analysis scripts available at DRYAD (datadryad.org), DOI:XXX.

## Abstract

Evolutionary theory is sufficiently well developed to allow for short-term prediction of evolutionary trajectories. In addition to the presence of heritable variation, prediction requires knowledge of the form of natural selection on relevant traits. While many studies estimate the form of natural selection, few examine the degree to which traits evolve in the predicted direction. In this study we examine the form of natural selection imposed by mantid predation on wing size and shape in the fruitfly, *Drosophila melanogaster*. We then evolve populations of *D. melanogaster* under predation pressure, and examine the extent to which wing size and shape have responded in the predicted direction. We demonstrate that wing form partially evolves along the predicted vector from selection, more so than for control lineages. Furthermore, we re-examined phenotypic selection after ∼30 generations of experimental evolution. We observed that the magnitude of selection on wing size and shape was diminished in populations evolving with mantid predators, while the direction of the selection vector differed from that of the ancestral population for shape. We discuss these findings in the context of the predictability of evolutionary responses, and the need for fully multivariate approaches.

## Introduction

Biologists measure natural selection to help identify agents of selection, to infer how current phenotypes were influenced by past selection and to predict future evolutionary outcomes. Since the publication of the seminal work by Lande and Arnold Lande and Arnold (1983), considerable effort has gone into measuring the form, magnitude and variability of phenotypic selection (Kingsolver et al., 2001; Hoekstra et al., 2001; Kingsolver and Diamond, 2011; Siepielski et al., 2009; Morrissey and Hadfield, 2012). However, additional factors influence the trajectory of evolution such as correlational selection, the stability of the selective function (Brodie III, 1992), as well as the genetic architecture of the traits (Hansen and Houle, 2008; Agrawal and Stinchcombe, 2009; Kirkpatrick, 2009), making such predictions difficult in practice. In this study we address this predictability, by investigating the extent to which experimental populations of *Drosophila melanogaster* subject to predation risk, evolve along the trajectory predicted from phenotypic selection.

Studies have investigated how closely populations evolve along the direction predicted from the multivariate breeders equation, (as a function of both selection and the genetic variance-covariance matrix) (Schluter, 1996; Hansen and Houle, 2008; Agrawal and Stinchcombe, 2009; Higgie and Blows, 2008; Hunt et al., 2007; Mcguigan et al., 2005; Walsh and Blows, 2009; Blows et al., 2004; Simonsen and Stinchcombe, 2010). Yet in only a handful of cases has selection been observed for a sufficient amount of time (beyond a few generations) to evaluate these predictions. Furthermore, the ecology and natural history of many organisms limits us to estimating phenotypic selection, generally over just a few generations (but see Ozgul et al., 2009; Grant and Grant, 2002, 2006). For some selective agents like predation, the organism is consumed, prohibiting (at least in the field) the measurement of many traits that are targets of selection. As a result, it may be challenging to predict the evolutionary trajectory of some traits involved with anti-predator activity. This might suggest a pessimistic view of our ability to predict the selective response in natural systems.

Despite these issues, convergent and parallel evolution are often observed among populations, suggesting that persistent and predictable selection may be relatively common (Conte et al., 2012), even if it is difficult to measure. While estimates of the strength of viability selection suggest it may be weaker than for other fitness components (Hoekstra et al., 2001; Lind and Cresswell, 2005; Ajie et al., 2007), repeated evolution of similar morphologies in response to predation for several fish species (O’Steen et al., 2002; Langerhans et al., 2004; Dayton et al., 2005; Langerhans and Makowicz, 2009) suggests a strong and consistent regime of selection. Similar results have also been observed for shell morphology among populations of snails in apparent response to predation (Auld and Relyea, 2011; DeWitt et al., 2000, 1999). When selection is relaxed by the removal of predators, even for just a few generations, trait means have been shown to change dramatically (Reznick et al., 1990, 1997; Reznick and Ghalambor, 2005), consistent with predation maintaining trait values in the face of potentially antagonistic selective effects. The prevalence of diverse, and often costly, traits that mediate interactions with predators suggests that predation profoundly influences fitness.

In this study we investigate how multivariate wing form of *Drosophila melanogaster* evolves along the trajectory predicted by initial estimates of phenotypic selection in response to predation by mantid nymphs (*Tenodera aridifolia sinensis*). This novel experimental system has a rather rare (but see Svensson and Friberg, 2007; Kuchta and Svensson, 2014) and useful attribute in which the wings are not consumed when the fly is captured by its mantid predator (figure 1A). This allows us to collect the wings from both surviving and consumed flies to estimate the form and magnitude of natural selection on both size and shape. Multivariate shape provides a robust framework for evaluating evolutionary predictions. While size only varies along one axis, a high dimensional representation of shape is less likely to change in the predicted direction by chance alone. This enables us to make clear quantitative comparisons of the degree of similarity between predicted and observed response to selection (Pitchers et al., 2013). It also extends a well developed genetic system for use in studies of predator-prey interactions.

**Figure 1.**
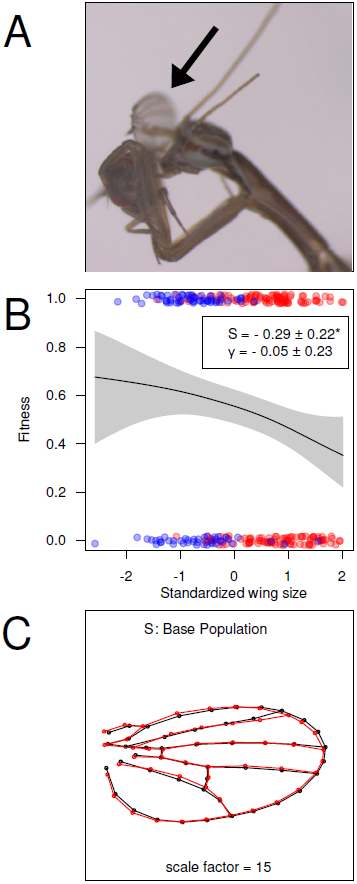
*(A)* 1st instar nymph of the Chinese mantid (*Tenodera aridifolia sinensis*) consuming a fruit fly. Note the wing about to drop off. *(B)* The selective function for size estimated by fitting cubic splines (*sensu* Schluter, 1988) along with estimates for linear and quadratic selection. Stars denote significance from logistic regression, but estimates are derived from a linear regression of size on relative fitness. Points above the function are individuals that survived. Points below the line were captured and eaten. Red dots are females and blue dots are males. Error bands are 95% confidence intervals. *(C)* Visualization of the selection differential (**S**) as measured in the base population. Points indicate landmarks and semi-landmarks. The shapes represent the mean shape plus 10x **S** (black line) and minus 10x **S** (red line).

Wing size and shape in *Drosophila* have been used as a model system for evolution (Gilchrist and Huey, 2004; Gilchrist et al., 2004; Huey et al., 2000; Gilchrist and Partridge, 1999; Weber, 1990*b*; Mezey and Houle, 2005; Pitchers et al., 2013), genetics, and development (Dworkin and Gibson, 2006; Houle and Fierst, 2013; Palsson and Gibson, 2000). There is substantial segregating variation for wing size and shape, with some variants mapped (Weber et al., 1999; Palsson, 2004; Zimmerman et al., 2000; Mezey et al., 2005; Mckechnie et al., 2010; Dworkin et al., 2005; Palsson et al., 2005). Studies have demonstrated that genetic variation is available along many dimensions of wing shape (Mezey et al., 2005). Using artificial selection, it has been demonstrated that this variation can be selected upon (Houle et al., 2003; Weber, 1990*b*, 1992, 1990*a*; Palenzona and Alicchio, 1973; Rochetta and Palenzona, 1975). Yet little is known about the selective agents influencing variation for wing form (but see (Hoffmann et al., 2007; Menezes et al., 2013)) or the potential functional role it plays in avoiding predation.

We quantified the magnitude and direction of selection on wing size and shape in an outbred population. We then allowed replicates derived from this population to experimentally evolve under episodic selection with mantids or under predator-free conditions. For the evolved populations we quantified changes in wing form, with particular focus on the direction of change, relating it to the vectors of phenotypic selection predicted from the base population. We demonstrate that while evolution of wing shape for the predator populations is more aligned with the initial vector of selection than are the controls, not all of the change is in the direction predicted by the initial vector of selection. Despite observing consistent directional selection on wing size, we observed considerable divergence in its evolutionary response. Furthermore, we measured phenotypic selection on the evolved populations, and demonstrate that the magnitude of selection on both size and shape is substantially diminished in the populations exposed to predation and is distinct from the initial vector of predicted selection. We discuss these results within the context of how populations change along a fitness surface, the importance of unmeasured traits and the degree of repeatability to agents of selection.

## Materials and Methods

### Base Populations

We used an advanced intercross with 100 inbred lines to generate a synthetic outbred ancestral population referred to as the base. The inbred lines were derived from two populations of wild *Drosophila melanogaster* collected in fruit orchards in Maine and North Carolina (Goering et al., 2009; Reed et al., 2010). Flies were round robin intercrossed for three generations and then allowed to mate randomly for 5 generations. We chose this approach, as a compromise to minimize confounding laboratory adaptation while still incorporating genetic variation present in natural populations. With this approach linkage disequilibrium among variants will likely be more extensive than in wild-caught flies. Post-intercross, we maintained the population at large size on cornmeal molasses media with live yeast in four 200ml culture bottles at 24°C and 60% humidity.

### Predation Environment

We used first instar nymphs of the Chinese mantid (*Tenodera aridifolia sinensis*) as predators. We collected mantid egg cases locally from old fields in southern Michigan and supplemented these from garden suppliers (Nature’s Control Medford, Oregon) when necessary. We hatched and maintained egg cases at 24°C and 60% humidity. Approximately 100–400 mantids emerged from each egg case and were used as predators for the duration of the first nymphal instar. After hatching, we housed mantids at 18°C and 60% humidity in arenas consisting of a 710mL plastic cup with a mesh covered window for air flow. We placed a tissue at the bottom of each cup to trap moisture when watering to help maintain humidity. We also added a green plastic aquarium plant to provide substrate for mantids to perch upon. Unless otherwise specified we used five mantids per arena.

All episodes of predation occurred at 18.5°C and 60% humidity, and were initiated between 12-3 pm. We fasted mantids for 24 hours before each episode of selection to increase predation rates. Arenas were cleaned with 70% ethanol and water before use. 25 flies were introduced into each arena via a funnel, after which arenas were returned to the incubator. After 24 hours, all predation arenas were moved to a 4°C refrigerator to knock down flies and mantids to aid collecting. We then removed the mantids from each container, and surviving flies were censused.

## Testing the role of flight in the escape response using a *vestigial^1^* mutant population

To test whether flight played a role in the escape response, we tested whether wing loss would negatively impact survival under risk of predation. We introgressed a mutation in the *vestigial* (*vg*^1^) gene into the base population by repeated backcrossing for 10 generations. The *vg*^1^ mutation causes a nearly complete loss of the wing blade and associated musculature (Sudarsan et al., 2001). We competed individuals from the *vg*^1^ mutant population with their wild-type conspecifics by placing 13 mutant and 13 wild-type flies in each of 16 arenas (8 arenas each for male and female flies). The survivors for both *vg*^1^ and their wild-type conspecifics were counted after 24 hours with the predators.

## Assaying Phenotypic Selection: Base population

To assess how naturally segregating variation for wing size and shape might be associated with survival during predation events, flies from the base population were exposed to predation. Predation on males and females was assayed in separate arenas so that we could examine independent effects of sex. We placed 20 flies into each arena (9 arenas each for females and males). Four days later we set up a second block of arenas (10 arenas for females, 8 for males). After predation, we collected all surviving flies and all wings from consumed flies from the bottom of the arenas. We also collected 100 individuals of each sex that were not exposed to predators. All bodies and wings were preserved in ethanol for dissection and measurement.

## Experimental Evolution

We randomly selected five hundred flies from the base population and used these as parents to generate the four populations for experimental evolution. We randomly assigned offspring of these 500 individuals to each of the treatments, with blocks of offspring for the different replicates. The predator free populations control for selection and adaption independent of the predators (i.e. alterations in competitive environment). Selection was administered in two replicate sets each consisting of one predation and one control population, hereafter referred to as PredR1, ConR1, PredR2, and ConR2. We offset the generational cycle of replicate 2 from replicate 1 by 2–5 days for logistical reasons, but the populations were otherwise treated identically. Each population was reared in four bottles with approximate densities of 100-500 eggs per bottle each generation. We did not explicitly control for density, but restricted egg laying time to 2–6 hours to avoid larval overcrowding. Bottles were reared at 24°C and 60% humidity until eclosion of adults.

Three days after eclosion of the first flies, progeny from each population were lightly anesthetized using CO_2_, placed randomly into vials, and maintained at 18.5°C and 60% relative humidity. The following day, flies from a given treatment were mixed under anesthesia, to minimize inadvertent selection on developmental time. Flies (25/vial) were transferred to fresh vials, corresponding to the number of arenas used for predation in that generation. Each generation we varied the total number of arenas depending on the voracity of the current batch of mantids in order to ensure that the total number of survivors was large enough to limit the effects of drift (between 150–400 surviving individuals/generation). Flies from the control vials were similarly mixed under anesthesia after which we placed 50 flies into each of 8 vials. Remaining predation and control flies were set aside as backups. All flies were then returned to the incubator at 18.5°C and 60% for at least 24 hours prior to the episode of predation. Because egg cases were seasonally available, we occasionally used second instar mantids to maintain experimental evolution. However, second instar mantids were not used for any experimental trials. Control arenas were identical to predation arenas, only lacking mantids.

After selection, we collected all survivors from the predation arenas. To maintain similar population sizes we selected individuals at random from the control populations matching the number of male and female survivors from the respective predation population. Individuals from each treatment were transferred into separate 30 × 30 × 30 cm polyester mesh cages, and allowed to recover for 30–45 minutes before fresh bottles of food media were placed into each cage. After allowing sufficient time for egg laying, the bottles were then removed from the cages and reared at 24°C and 60% humidity. After breeding, remaining adult flies were stored in ethanol at −20°C.

## Assaying Phenotypic Selection: Evolved populations

To examine how the fitness function changed as a result of experimental evolution, we repeated phenotypic selection (as described above) during generations 31 and 32 of the experiment. Given the large size of this experiment, it was performed in four blocks, with two blocks for each generation. At generation 31 of experimental evolution, we set up 14 arenas each of PredR1 females & males and ConR1 females & males. Five days later, we set up 14, 14, 8, and 9 arenas for PredR2 females & males and ConR2 females & males respectively. At generation 32, we set up 14 arenas each of PredR1 females & males and ConR1 females & males. Five days later we set up 13, 13, 14, and 14 arenas for PredR2 females & males and ConR2 females & males respectively. As before, we collected all surviving flies and unconsumed wings and stored them in ethanol. Overlapping egg cases were used for this experiment, and egg case of origin was used as a covariate in the model (see below). We distributed mantids so as not to confound predation effects across replicates and treatments.

## Wing Measurement & Statistical Analysis

Wings were dissected and mounted on slides in 70% glycerol. When available, both wings from an individual were mounted. Wings were also dissected from 25 flies that were stored from the initial generation of experimental evolution, and from every 10 generations following up to generation 30 to estimate the trajectory of size and shape change. Wings were imaged at 40X magnification on an Olympus DP30BW camera mounted on a Olympus BX51 microscope using ‘DP controller’ V3.1.1 software. All images were saved in greyscale as tiff files.

To capture landmark and semi-landmark data we followed a modified protocol (Pitchers et al., 2013) for the use of the wingmachine software (Houle et al., 2003). We used the program tpsDig2 (Rohlf 2011) to manually record the coordinates of two starting landmarks, and used wingmachine to fit nine B-splines to the veins and margins of the wings in the images. We extracted 14 landmark and 34 semi-landmark positions, and performed Procrustes superimposition (Zelditch et al., 2012). After superimposition, the positions of semi-landmarks were allowed to slide along each segment of the wing margin/veins, minimizing Procrustes distance, using CPR V0.2 (Marquez 2011). The data were checked for visual outliers at multiple stages; and putative outlier images were reexamined and splines re-fit if necessary. The (semi-)landmark configurations for all wings measured for this study were superimposed together, resulting in a common shape space. Centroid size was used as measure for size (Zelditch et al., 2012). For flies with both wings collected, we calculated the mean shape and centroid size per individual.

## Model Selection

For the univariate analyses, we evaluated model fits using Akaike’s Information Criteria (AIC) and Bayesian information criteria (BIC). AIC has been shown to often ‘prefer’ more complex models than there is support for, particularly when sample sizes are large (Grueber et al., 2011), we used model weights from BIC throughout for consistency to perform model averaging when appropriate. Unless otherwise noted, all further analyses were conducted in R (V2.15.1) (R Core Team, 2012). Scripts for the custom functions described below are available with the data from DRYAD.doi.XXXX.

## Analysis of Survival

For the *vg^1^* mutant and wild-type competition assays, we fit the model:

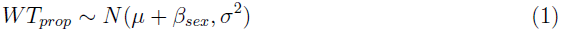

where *WT_prop_* was the proportion of wild-type survivors in each arena, and *β_sex_* was the model coefficient for sex.

For the base and evolved populations, we measured survival ability as the total number of surviving flies in each container. For the base population, we fit the model:

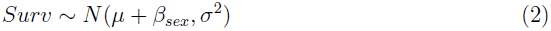

along with a set of expanded and restricted models (Supplementary table 1) where *Surv* was the number of surviving flies in each arena and *β_sex_* was the model coefficient for sex. Model averaging produced coefficient estimates indistinguishable from the model with best support so this model was used for further inference.

For the evolved populations we fit the model:

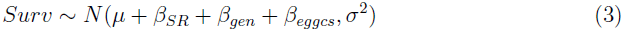

along with a set of expanded and restricted models (supplementary table 2) where *Surv* was the number of surviving flies in each arena and *β_SR_*, *β_gen_*, and *β_eggcs_* were the model coefficients for selection regime, generation of selection when the assays were performed, and the egg case of origin for the mantids in each arena. The coefficient estimates produced from model averaging models 1 and 2, which accounted for 95% of the overall support, were indistinguishable from the model with best support so for simplicity we used it for further inference. For the above models, we confirmed the effects using generalized linear models (poisson with log link), or a logistic regression with similar results.

## Analysis of Phenotypic Selection on Size

We used the Lande and Arnold (1983) approach to examine selection acting on wing size in the base and evolved populations. As recommended by Janzen and Stern (1998), we used logistic regression on survival for statistical inference and general linear model on relative fitness to estimate coefficients. Relative fitness for each individual was calculated by scaling survival (0 for dead and 1 for survived) by the total proportion of survivors in each experiment. To measure linear selection in the base population we fit the model:

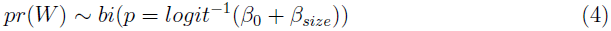

along with a set of expanded and restricted models (supplementary table 3). *W* was absolute fitness (survival) and *β_size_* was the model coefficient for standardized wing centroid size. The coefficients produced by model averaging were indistinguishable from the model with best support so it was used for further inference. It should be noted that we are estimating the linear S, and non-linear *C* selection differentials (Brodie III et al., 1995). The *β*’s in the equations are used to represent estimated model parameters, and do not represent selection gradients.

To measure linear selection in the evolved populations we fit the model:

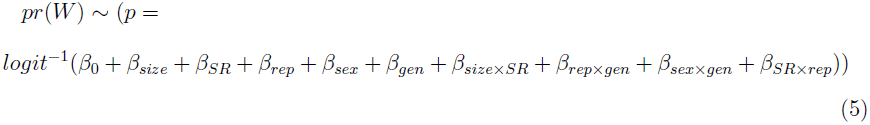

along with a set of expanded and restricted models (supplementary table 4) where *W* was fitness and *β_size_*, *β_SR_*, *β_rep_*, *β_sex_*, and *β_gen_* were model coefficients for standardized wing centroid size, selection regime, replicate, sex, and the generation of experimental evolution respectively. We fit separate models as above including the quadratic effect of size to estimate non-linear selection. Estimates for non-linear selection on size were near zero, non-significant, and did not improve model fits in either the base or the evolved populations.

Non-parametric estimation of the form of the fitness functions substantially aids visualization and interpretation of fitness functions (Schluter, 1988). We therefore used generalized additive models from the the mgcv package V1.7.22 (Wood, 2004) to fit cubic splines to subsets of the data from each experiment corresponding to the relevant significant effects estimated by the logistic regression analyses. Optimal smoothing parameters were estimated using REML.

### Variances in Size & Shape

One additional approach to investigating natural selection is to examine the changes in phenotypic variance before and after the selective event (Endler, 1986). Under either directional or stabilizing selection, a reduction in variation would be predicted. Under disruptive selection however, we would predict an increase in variation. Analyzing differences in variance between dead and surviving flies, as well as between predation and control populations may therefore provide additional information on the type of selection occurring in these populations. We used Levene’s test to assess changes in variance for wing size, using deviations from the median rather than the mean since this approach is more robust to departures from normality. For the base population, we modeled the main effects of sex and size because our previous analyses lacked support for an interaction between sex and the form of selection. We fit the model:

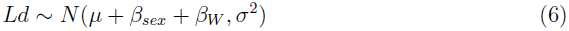

where *Ld* was the Levene’s deviates for each individual, *β_sex_* was the model coefficient for sex and *β_W_* is the model coefficient for absolute fitness. Though our previous analyses do not suggest that selection acting in the evolved populations differed between replicates, differences in size between PredR1 and PredR2 suggest that its inclusion is appropriate. We fit the model:

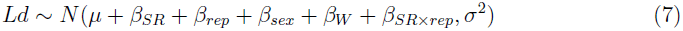

where *Ld* was the Levene’s deviates for each individual and *β_SR_*, *β_rep_*, *β_sex_*, and *β_W_* were the model coefficients for selection regime, replicate, sex, and absolute fitness, respectively. Confidence intervals for all estimates were generated by non-parametric bootstraps, in order to avoid issues with non-normality of residuals. We also calculated the coefficient of variation for each of the groups modeled above. Because we know that there is substantial sexual size dimorphism as well as size differences among the base and evolved populations, this approach should provide more intuitive visualization.

To compare levels of variation in shape we took a somewhat simpler approach. We expressed the variability of each group as the trace of its covariance matrix for shape. We then bootstrapped the data to generate samples of each covariance matrix in order calculate confidence intervals on the estimated matrix trace. Non-overlapping (95%) confidence intervals intervals were then used to infer statistical support for differences in variance among groups.

### Multivariate Analysis of Shape

In our initial analyses, we found that the modeled effects of allometry and sexual dimorphism were extremely consistent between treatments and over time (i.e. the vectors of model coefficients for sex and for size were very tightly correlated; see below). In order to facilitate the interpretation of the modeled coefficients of selection and generation number, we therefore sought to exclude these effects from our analyses. With data from all wings pooled, we fit the model:

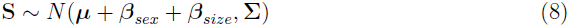

where **S** is the matrix of Procrustes coordinates and ***β****_sex_* and ***β****_size_* are the vectors of model coefficients for sex and for wing centroid size respectively. Σ is the “error” covariance matrix. We retained the residuals from this model as our shape variables.

Configurations of Procrustes coordinates by definition include dimensions without variance. The Procrustes superimposition results in a deficiency of 4 ranks (1 each for removed size and rotation information, and 2 for position), and each semi-landmark may contribute as little as 1 added dimension (Zelditch et al., 2012). In order that the shape data would not be rank deficient, we extracted principal components from the (96-dimensional) residuals, and retained the first 96 − (4 + 34) = 58 principal components, comprising > 99.9% of the shape variance in the full set of residuals. Shape PC’s used in all the analyses below are thus of full rank, and are expressed in a common sub-space.

## Modelling Shape Change

Separately within each of the four evolved populations, we estimated the direction of observed evolutionary change as the vector of model coefficients from the multivariate linear model:

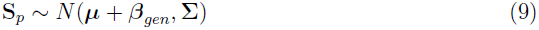

where **S***_p_* is the matrix of principal component scores for shape in a given population and ***β****_gen_* is vector of model coefficients for time, expressed as the number of generations removed from to the base population. Once we had estimated these vectors of parameters, we compared their directions by calculating vector correlations as:

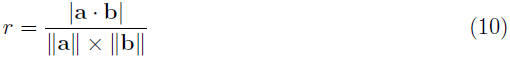

where |**a** · **b**| is the absolute value of the dot (scalar) product between vectors **a** and **b**, while ||**a**||, and ||**b**|| are the magnitudes (L^2^, or Euclidean norm), for each vector. The absolute value for the dot product was used to avoid any numeric issues with arbitrary sign changes that can occur computationally (during the bootstrapping procedure, see below). Thus *r* = 0 represents no similarity between the vectors while *r* = 1 means the two vectors point in an identical orientation (but possibly opposite in direction). Given that *r* is a multivariate extension of the Pearson correlation co-efficient *ρ*, we consider this a more intuitive measure than the vector angle (*θ* = arccos (*r*) in radians) which has been used elsewhere. Confidence intervals were computed using non-parametric random pairs bootstrapping, from 10,000 bootstrap iterations. This approach was used both to compare the direction of **S** as measured in all five populations, and to compare the directions of observed shape change among the evolved populations.

To illustrate the magnitude of change in wing shape during experimental evolution, we calculated a shape score (Drake and Klingenberg, 2008). Briefly, we projected the shape data onto a line in the direction defined by the vector of model coefficients for the generation term (***β**_gen_*) from model (3):

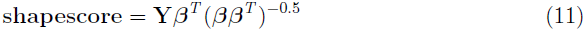

The shape score provides a univariate measure of shape change that can be plotted against generation number to visually assess the magnitude and linearity of the relationship (Drake and Klingenberg, 2008). We used custom R functions to calculate vector correlations and shape scores.

### Selection on Shape

Within each population, we estimated the vector of linear shape differentials (**S**). Traditionally this would be calculated as vector of differences between the mean phenotype of survivors and the mean phenotype of those individuals that were preyed upon. Here we estimated **S** using a 2-block partial least squares (PLS) approach (Rohlf and Corti, 2000; Klingenberg and Zaklan, 2000; Mitteroecker and Bookstein, 2011; Klingenberg and Monteiro, 2005; Gomez et al., 2008, 2006) with the matrix of the 58 shape PC’s forming one block, and the vector of survival data (0 or 1 for dead or survived) as the second block. We note that in this case this estimate of **S** is proportional to the differences between the mean shape configurations for the dead and survivors.

It is important to note that wing shape itself is the trait, and not individual landmarks/PC’s. After Procrustes superimposition, individual landmarks and semi-landmarks cannot be meaningfully interpreted independent of the whole shape configuration and the superimposition can generate correlation between landmarks that is confounded with biological correlations (Zelditch et al., 2012). Thus, interpreting the selection gradients, ***β***, from a multiple regression for shape (*sensu* Lande and Arnold, 1983) for “individual” shape variables is biologically meaningless (Albert et al., 2008). In addition, selection gradients can be difficult to visualize for shape (Klingenberg and Monteiro, 2005) but see (Mitteroecker and Bookstein, 2011), in particular because estimating the inverse of the phenotypic covariance matrix, **P**^−1^, can be problematic. We observed that, upon resampling, lack of stability in **P**^−1^ caused computationally difficulties. One alternative is to retain only the first few PC’s and analyze them as if they were independent traits (Gomez et al., 2006, 2008; Kuchta and Svensson, 2014). This is still sub-optimal, however, since substantial variation and selection may be missed and the biological interpretation of any selection that is detected is difficult. While this is an important and outstanding issue, we elected to use selection differentials for the shape analyses because they retain biological meaning and the focus of the study is on the predictability of selection not its specific form or estimation. However, this does mean that the results need to be interpreted as a combination of both direct and indirect selection on shape.

We estimated total selection on wing shape as the magnitude of the vector of the selection differentials, ||**S**||, and used sampling with replacement of the data to generate non-parametric bootstrap confidence intervals on these estimates. Additionally, we permuted survivorship relative to the measures of shape to assess the null hypothesis that wing shape does not contribute to variation for survivorship. We also compared the directions of the **S** vectors using vector correlations as described above. Finally, we wanted to assess the degree to which the experimental evolution populations had evolved in the direction ‘predicted’ by selection as measured in the base population. To do this we calculated the vector correlations between the **S** vector measured in the precursor population and the vector of model coefficients for generation (***β****_gen_* from model (8)) as modeled separately for each population.

## Results

### Evidence that flight aids in the escape response

To test whether flight performance and wing form were potential targets of selection driven by the mantid predators, we introgressed a mutation in the *vestigial* (*vg)* gene into our base outbred population that nearly completely ablates the wing blade and associated flight muscles. We competed *vg*^1^ (functionally wingless) flies and their wild-type conspecifics with the predators. As predicted, the *vg*^1^ individuals were disproportionately the targets of predation. The survivors for both sexes consisted of approximately 60% wild-type and 40% mutant individuals (figure S1), consistent with a role for flight and possibly wing morphology in the escape response of *Drosophila*.

### Predator driven selection on natural variation for wing form

We next asked how natural variation for wing form was associated with survivorship by exposing flies from the base population to the mantids. We observed evidence for significant negative directional selection on wing size (Figure 1B, Table S1, S = −0.29 ±0.22, p ≃ 0.01) with little evidence for non-linear selection (*C* = −0.05 ±0.22, p ≃ 0.16). Visualization by fitting cubic splines to the survival data (Schluter 1988) was consistent with the estimates of directional selection (figure 1B). Despite sex specific differences in survivorship (6.7 ±0.75 & 3.8 ±1.07 survivors per arena for females and males respectively), evidence was weak for an interaction between selection on size and sex (Table S1). It is currently unclear whether the target is wing size *per se* or whether it is due to a correlation between wing size and overall body size.

Additionally, shape has been shown to be correlated with escape ability in other organisms (Langerhans et al., 2004; Dayton et al., 2005; Langerhans and Makowicz, 2009; Svensson and Friberg, 2007). Using a 58 dimensional representation of wing shape (figure S2), we used partial least squares (PLS) to estimate the vector of selection differentials for shape (**S**) in the base population. We visualized this **S** vector for shape comparison (figure 1C). We see selection for a change in aspect ratio in which relatively longer and narrower wings are favored.

### What does wing form look like after experimental evolution?

We asked whether experimental evolution under risk of mantid predation would result in phenotypic changes in wing form consistent with our initial estimates of selection. Previous work has demonstrated that both wing size and shape have moderate to high heritabilities, and most aspects of wing shape are readily altered by artificial selection (Mezey and Houle, 2005; Weber, 1990*b*). To assess this we subjected two populations of flies to episodic selection by mantids, each paired with control populations evolved without predators. The control populations allow us to assess the effects of selection due to the experimental evolution procedure unrelated to the predators that occurred during the experimental evolution process.

As expected, survival in both predator populations increased, compared to the controls (10.86 ±0.78 & 13.98 ±0.79 survivors per arena for control and predation populations respectively; figure 2A). This represents a ∼30% increase in survivorship relative to the control populations. We did not observe differential survival between sexes in this experiment for either selection regime.

**Figure 2.**
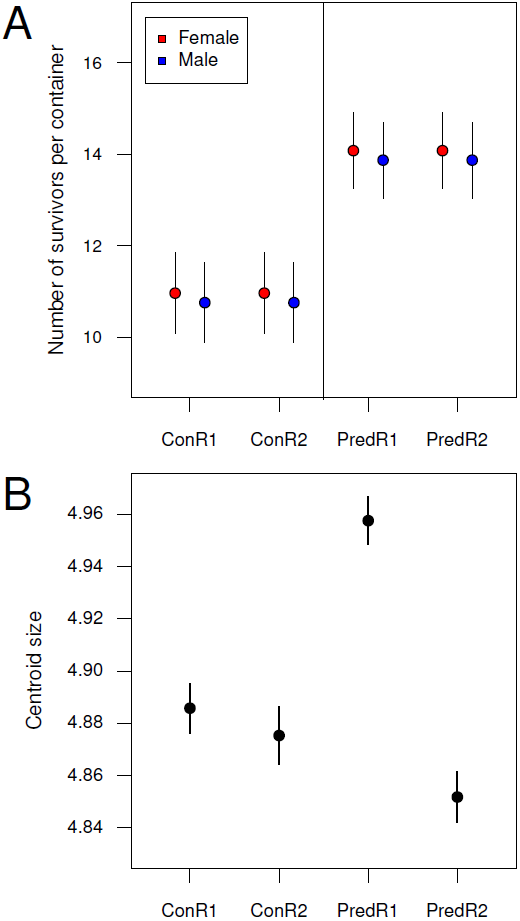
*(A)* Mean number of survivors in each predation arena after 30 generations of experimental evolution. *(B)* Differences in wing size of the evolved populations after 30 generations of experimental evolution. Female values shown are 13% larger than those for males. Errors are 95% confidence intervals.

We measured individuals stored during the experimental evolutionary process every ten generations, from the base to generation 50, to track changes in wing form. Wing size of all populations increased ∼3.7% over the 50 generations of experimental evolution (0.0035 mm ±0.0004 mm per generation). This change was most likely a result of selection due to non-predatory aspects of the experimental evolutionary procedure because all populations increased at similar rates and maintained the same overall sexual dimorphism. However, environmental variation in these samples collected directly from the experimental evolution regime is relatively large (Figure S3).

To more carefully estimate size differences among the evolved populations, we measured wing size in the overall population by using the wings from the dead and surviving flies from the phenotypic selection experiments, as all flies were reared under density controlled conditions. Thus environmental and genetic effects were not confounded. Comparison of the number of surviving and dead flies from this assay to the number of wings recovered suggests that nearly all wings from dead individuals were recovered, and should provide reasonable estimates.

Under these conditions, the relevant contrast is the difference between the control populations and predation populations. We found that the two control populations had similar wing sizes, yet the two populations evolved under risk of predation diverged in size (figure 2B) even though all populations showed a general size increase relative to the ancestral population. Surprisingly only PredR2 has evolved in the predicted direction, with wings ∼1% smaller than controls, while PredR1 evolved wings that are ∼2% larger.

In terms of shape, all four populations have evolved from the base population, though not to an equal extent. We visualized the evolutionary trajectories of the four populations by plotting the shape score for generational effects (equation 10) (figure 3A & B). In all four cases the evolutionary trajectories were best described by a simple linear model. Whereas the two control populations have changed in a very similar fashion (figure 3A), the two predation populations are clearly divergent, with wing shape in PredR2 evolving significantly more rapidly than in PredR1 (figure 3B).

**Figure 3.**
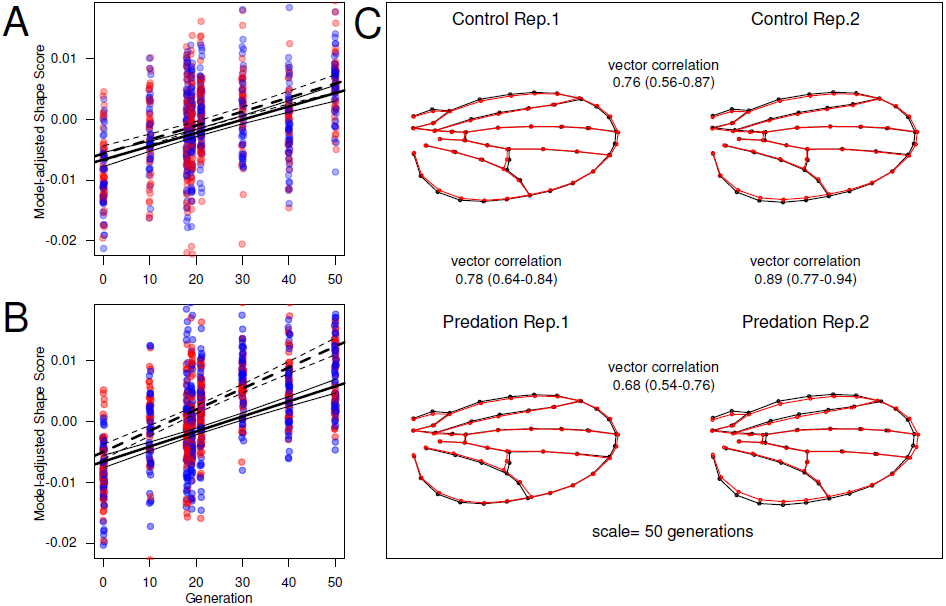
*(A)* Shape score by generation for control *(B)* and predation selection regimes. Model adjusted shape score for generation (*sensu* Drake and Klingenberg, 2008 see methods) is plotted against generation number, with blue points for males and red points for females. Solid regression lines and 95% confidence intervals are for replicate 1, and dashed lines and 95% confidence intervals are for replicate 2. *(C)* Visualization of the directions of the evolution of wing shape in the 4 experimental evolution populations. The shapes represent the mean plus (black line) and minus (red line) the modelled vector of evolutionary change in each case, scaled to 50 generations in magnitude. The points represent landmarks and semi-landmarks. Vector correlations between these modelled directions of shape evolutions (and their 95% credible intervals) are printed between the pairs of populations to which they relate.

Over the course of experimental evolution the wings of all populations have changed aspect ratio: their length increasing slightly as their depth decreases. This change is most pronounced in PredR2 (figure 3C). Other than the differences in aspect ratio, PredR1 and PredR2 differ most noticeably in the response of the cross-veins and the distal end of L5. PredR1 demonstrates a proximal shift in the posterior cross-vein and an anterior shift in the attachment of L5 to the margin; by contrast in PredR2 there was no change in L5 and an anterior shift in the anterior cross-vein.

### What do the fitness functions look like after experimental evolution

After 30 generations of experimental evolution, we again exposed flies that evolved with (and without) predators to a bout of predation. We observed negative directional selection for size in the control populations (S = −0.16 ±0.07, p ≃ 0.0001; figure 4A), consistent with our findings from the ancestral base population. We also observed negative directional selection in the predation populations, but of diminished magnitude relative to the controls (S = −0.06 ±0.05, p ≃ 0.0005; figure 4B). Both the control and the predation populations showed extremely weak quadratic selection on size (*C* = 0.0001 ±0.03, p ≃ 0.73). We might reasonably expect that the reduction in the magnitude of directional selection in PredR2 was a result of evolutionary change in wing size in response to selection. However, this provides no explanation for the increase in size in PredR1.

**Figure 4.**
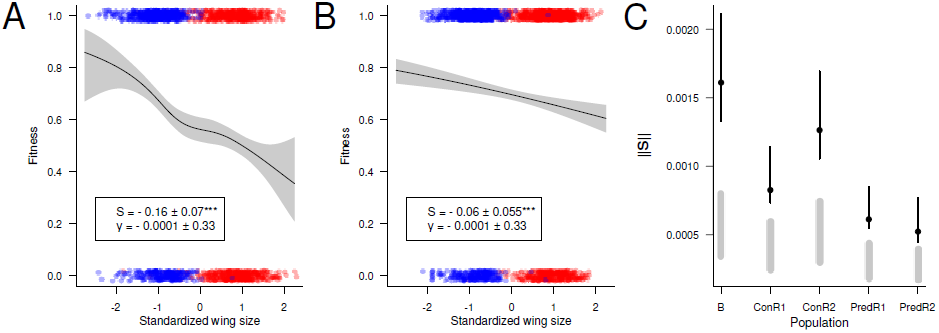
The selective function for size estimated by fitting cubic splines (*sensu* Schluter, 1988) with replicates pooled for the *(A)* control *(B)* and predation populations along with estimates for linear and quadratic selection. Points above the function are individuals that survived. Points below the line were captured and eaten. Red dots are females and blue dots are males. Error bands are 95% confidence intervals. *(C)* Magnitude of the selection differential (**S**) as measured in the base (B), the control, and predation populations. Black points and lines are estimates and bootstrapped 95% confidence interval. The grey lines are the 95% confidence intervals from permutation of the same data; they represent the null hypothesis that the magnitude of **S** is random relative to survival.

We assessed selection on wing shape in the evolved populations as **S**; the vector of selection differentials between captured flies and survivors and compared the magnitudes of total selection on shape from the differential, ||S||. For the estimates of ||S||, we also generated distributions under the null expectation (of no association between wing shape and survival) using permutations of the data. In addition we also calculated the vector correlations between differentials in order to quantify their degree of alignment. For both approaches we computed confidence intervals on our estimates by applying a non-parametric bootstrap approach.

As can be seen from figure 4C, there was evidence for a significant association between shape and survival in the presence of the mantid predators for all populations: the estimates of ||S|| exceeded the 95% threshold permuted under the null hypothesis. Also notable are the much smaller ||S|| estimates of the predation populations compared to that in the base population. This evidence is consistent with a relative reduction in the magnitude of selection experienced by the predation populations after 30 generations. Interestingly, there is some evidence for difference in ||S|| between the two control populations, however both still exceed the predation treatment regimes.

The directions of the **S** vectors in the predation populations are also quite divergent, however, and their vector correlations with the base differ; 0.39 (0.07–0.59) & 0.02 (∼0 – 0.34) though our estimate are not very precise (vector correlation followed by bootstrapped 95% confidence intervals, base **S** vs. PredR1 **S** & PredR2 **S** respectively). By contrast the vector correlations between the controls and the base population were consistent with one another at 0.39 (0.01 – 0.51) & 0.29 (0.04 – 0.48) respectively (see figure S4 for all pairwise comparisons). Moreover, the directions of the **S** vectors measured in the four evolved populations are not closely aligned (figure S5).

### Has wing form evolved in the direction predicted by selection on the base population?

While there has been evolution of shape in all four populations, we wanted to assess how much of the observed change is in the predicted direction (based on **S** in the ancestral base population). We calculated the vector correlations between the generation shape change vectors from each evolved population and the **S** vector from the base population and observed that the evolutionary responses of the predation populations were more aligned with the predicted vector compared with the control populations (figure 5). It is notable that none of the populations are particularly highly aligned with the initial predicted vector.

**Figure 5.**
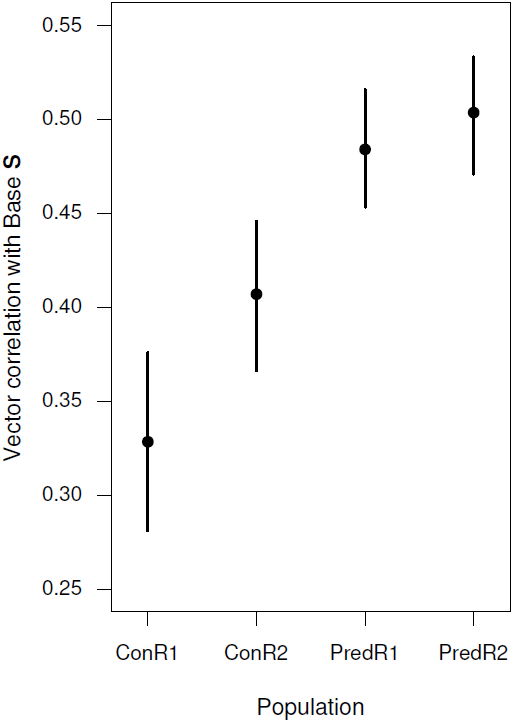
Vector correlations between **S** for wing shape estimated in the base population, and the direction of the direction of shape change during experimental evolution. The response vector was estimated within each population. Points are vector correlation estimates, and lines represent 95% bootstrapped confidence intervals.

Given that both predation populations have experienced a similar reduction in the magnitude of selection (as represented by ||S||: figure 4C), and a similar amount of evolutionary change in this direction (figure 5), it appears that they experienced similar changes in the selective function for shape, despite their divergent evolutionary response for size (figure 3C). This suggests that the two predation populations are evolving different avoidance strategies in response to the predation pressure imposed by the mantids — likely involving traits other than wing morphology. In the case of PredR2 there is evidence that the reduction in the intensity of selection is associated with evolution in the predicted direction, but it seems likely that there may be other adaptions occurring in PredR1

### Changes in variance in both size and shape

Changes in variation can also provide information about the form of selection experienced by a population. Evidence for differences in variance for size between survival classes in the base population was weak (Levene’s deviates = 0.28 ±0.04, 0.27 ±0.03, 0.24 ±0.04 for unselected, dead, and surviving individuals respectively, p ≃ 0.25). Males had lower variance for all survial classes (−0.07 ±0.03, p < 0.001). For the evolved populations, differences in size variance were only found among populations, with ConR1 and PredR2 having equal variance (Levene’s deviates = 0.11 ±0.01, 0.11 ±0.01 respectively), ConR2 having higher variance (Levene’s deviates = 0.15 ±0.01), and PredR1 having lower variance (Levene’s deviates = 0.10 ±0.01). Surviving individuals trended towards lower variance, but the difference was not significant (−0.003 ±0.006, p ≃ 0.37). Males again had lower size variance, but a much lower magnitude of difference (−0.015 ±0.006, p < 0.005).

Estimates of variance for shape in each population (the trace of the covariance matrix) show a dramatic reduction in variance in surviving flies and lower overall shape variance in the populations that evolved under predation risk (figure 6). Not only is the variance lower in the predation populations as compared to the controls for shape, but the surviving populations have much lower variation when compared to the populations that were captured and eaten by the mantids suggesting that selection has already reduced variation in the evolved populations and continues to do so.

**Figure 6.**
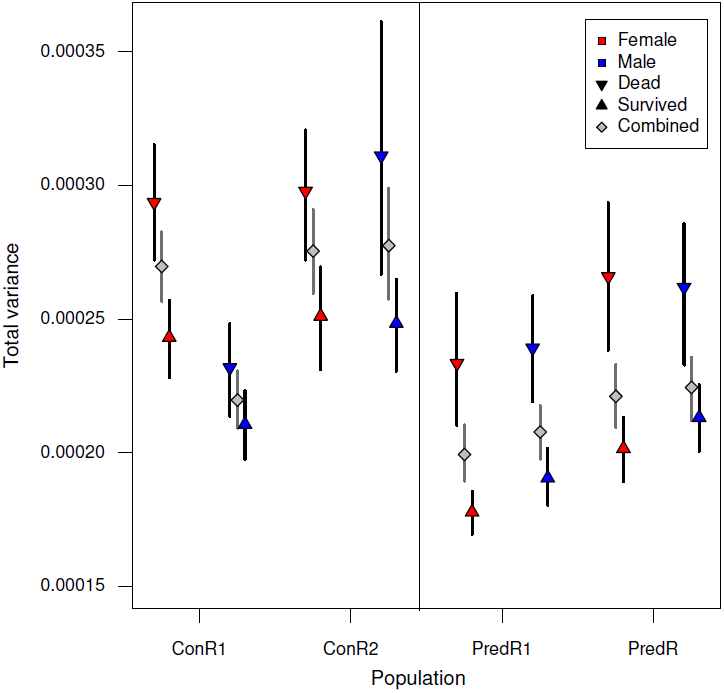
Estimates of variance for shape calculated as the trace of the covariance matrix for female and male flies from the evolved populations. Estimates of total variance (grey diamonds) are calculated with dead and surviving flies combined. Error bars are 95% bootstrapped confidence intervals.

## Discussion

For several decades, phenotypic selection analysis has been used to attempt to identify the primary targets of selection within natural populations under the assumption that the presence of selection on specific traits would provide information about about how those traits evolved and what future changes could be expected. The striking levels of convergence and parallellism in several well known study systems suggests that this assumption may be valid (O’Steen et al., 2002; Langerhans et al., 2004; Dayton et al., 2005; Langerhans and Makowicz, 2009; Auld and Relyea, 2011; DeWitt et al., 2000, 1999). However estimates of phenotypic selection analyses have not always been able to predict long-term evolution. It remains unclear whether this contradiction is a result of publication bias or indicative of larger issues. In this study we measured phenotypic selection on a naive population in response to a novel predator. We then re-measured the strength and direction of natural selection after populations were allowed to evolve under natural selection with the predator to determine whether our results would match the patterns of parallel response cited above. We found that the populations evolved divergent morphology for size but relatively consistent shapes. What do these results tell us about the form of natural selection, and what are the implications for its use in evolutionary prediction?

To use the breeder’s equation to predict an evolutionary response, we require not only a vector of directional selection, but heritable variation along the same axis as selection (Hine et al., 2011; Walsh and Blows, 2009). The direction of this genetic variation is determined by the size and structure of the genetic covariance matrix **G**. A number of studies have demonstrated that populations tend to evolve along genetic lines of least resistance (Schluter, 1996; Mcguigan et al., 2005), not necessarily the direction of strongest selection.

Previous work has demonstrated that there is considerable segregating genetic variation in most populations for wing shape. In particular the effective dimensionality of **G** for wing shape is quite high, and close to the number of measured traits (Mezey and Houle, 2005). Given the size and scope of the endeavor, we did not attempt to estimate **G** for the base population we used. However, genetic variation among the progenitor strains used to generate the population shows a high effective dimensionality (data not shown), consistent with previous results from other populations. It is possible that genetic variation in the direction of selection imposed by mantid predation may be minimal, and that the genetic line of least resistance is not perfectly aligned with this direction. Thus at least some of the common changes in wing form may be the result of a combination of lab domestication and evolution along the genetic lines of least resistance. Despite this, we see clear evidence for more shape change in the predation regimes consistent with the initial vector of selection. Given the high dimensionality (58) of shape, this is a pronounced effect, demonstrating that even with potentially countervailing selective and genetic forces, selection is still altering shape as predicted.

In addition to the need for available genetic (co)variation, there are several factors that can influence the evolutionary response to directional selection including: indirect selection due to correlated traits, stabilizing selection, fluctuating selection, fitness trade-offs, and unmeasured (but strongly selected) traits (Kingsolver and Diamond, 2011). The selection differentials reported include direct and indirect selection, so even though we cannot estimate the separate contributions of each, we saw significant total selection on both wing size and shape. Additionally, neither the base nor the evolved populations showed evidence for stabilizing selection that might have reduced the strength of directional selection. Though we expected the base population to be far from any fitness peak, it is curious that the predation populations did not show stronger non-linear selection given the reduction in the strength of the linear component.

## How similar is the form of selection after 30 generations of experimental evolution

The selective pressure imposed by the mantids on size remained relatively stable, consistent with the conclusions reached by Morrissey and Hadfield (2012). Our measure of S in the ancestral base suggested strong directional selection for smaller wings (∼−0.29). After nearly two years and 30 generations of evolution, the estimate of S was remarkably similar in the control populations (−0.16, figure 4). This slight reduction in the magnitude is, perhaps, unsurprising since the difference in sample size—nearly an order of magnitude greater for the evolved populations—allowed for more precise estimation. The estimate is also in line with the median reported by Kingsolver and Diamond (2011) for size traits (|0.14|) and viability (|0.08|). As a result, there is little evidence to suggest that temporal variation in selection from generation to generation resulted in the divergence in size in the predation populations particularly because both populations still exhibit measurable directional selection.

### Selection on shape

For shape, the picture is less clear. Because of the high dimensionality of shape, not only is estimation much more difficult, but there is a much larger available phenotype space. Perhaps unsurprisingly then, the vector correlations between selection in the base population and the control populations are reasonably low (∼0.35) suggesting that fluctuating directionality of selection may have limited the evolutionary response. The degree to which there was true variation in direction of selection for shape, as compared to estimation issues (even with our large sample sizes) remains unclear. Indeed, this is one of the major reasons we used **S** instead of *β*, as estimating **P**^−1^ proved to be computationally difficult, and caused problems during resampling. Despite this, both PredR1 and PredR2 show considerable overlap between the vector of shape change during evolution and the direction of selection predicted in the base population (*r* ∼0.5, figure 5). This suggests that even though the form of selection for shape is apparently less stable than for size, it has not resulted in substantial divergence between populations.

It is worth considering what is lost by using the selection differential **S** instead of the gradient, *β* = **P**^−1^**S**. For most phenotypic selection studies the main difference relates to disentangling direct and indirect selection on traits (pre-multipling by **P**^−1^ removes the phenotypic covariation). Shape data is unique, in that the different variables are not independent traits. Instead the whole configuration (as represented by a vector for each individual) is a geometric representation of the shape “trait”. Pre-multiplication by **P**^−1^ has the potential to change the observed orientation of the vector of the selection differential **S**, however it also causes difficulties with interpretation of the resulting selection gradients, and so the preferred method is to visualize the selection differentials (Klingenberg and Monteiro, 2005) as we have done here (but see (Mitteroecker and Bookstein, 2011) for an alternative perspective). Other groups have instead utilized a small number of principal components of the shape data in a standard Lande–Arnold selection gradient analysis (Gomez et al., 2006; Kuchta and Svensson, 2014). However, this utilizes a fraction of the variation in shape, with no guarantee that it represents the components of variation under selection. Thus a full multivariate approach is needed (Klingenberg, 2010) though we currently lack an accepted standard *sensu* Lande-Arnold. We suggest that continued effort and discussion into estimating and visualizing selection on shape, as well as determining the appropriate “dimensionality” of such effects is warranted.

### Possible causes of divergence and parallel evolution

Unknown fitness trade-offs and lab adaptation may have played a role in the divergence between PredR1 and PredR2. During the course of evolution, all four evolved populations showed a net increase in wing size of ∼3.2% and a lengthening and broadening of the wing blade in direct contrast to the smaller, longer, and narrower wings favored by selection in the base population (figure 1). These changes were remarkably consistent among the four populations and are likely a result of selection due to shared aspects of the experimental evolutionary process independent of the predators. However, though the directionality of the shared evolved response and selection measured in the base population suggests that evolution may be slowed in the predation populations, this gives little indication as to why PredR1 diverged from the predicted size trajectory. This could be caused by drift between the replicates over the 30 generations of experimental evolution. However, this explanation is unsatisfactory as the evidence of drift is missing in the control populations wherein it would be expected to dominate. Though the utmost care was taken to control variation between the replicates, differences in the health and voracity of the mantids, as well as in some other environmental factors was unavoidable, possibly contributing to this effect.

Where do these results leave us? We possess robust theory for measuring selection, and for predicting evolutionary responses into the near future (Lande and Arnold, 1983). However, we are often left to assume that populations will evolve phenotypes in the distant future consistent with these estimates. Though a number of other systems have examined the evolutionary consequences of manipulating predation regimes long term, notably the work of David Reznick and colleagues (Reznick et al., 1990, 1997; Reznick and Ghalambor, 2005), few studies have investigated how well evolutionary responses coincide with specific measures of selection. The *Drosophila*-mantid system described here allows us to maintain specific selective pressure in a relatively homogeneous environment on a population with a known history. This allows us to not only impose specific selection pressures, but to remeasure selection itself during the evolutionary process.

It is likely that unmeasured anti-predator behavioral traits played an important role. Unmeasured traits that may be under selection (and genetically covary with measured traits) can profoundly influence the biological inferences we make about natural selection, and evolutionary response. While many studies of phenotypic selection attempt to examine multiple traits that mediate the ecological interactions that generate variation in fitness, it is impossible to capture all of them in any one study. In a system like ours, where we employed a novel predator for *Drosophila*, anti-predator behaviors that were initially rare in the progenitor population can rise in frequency, fundamentally changing aspects of selection on other traits. For instance, if the escape response to direct attacks was the primary strategy early in the evolutionary process, but the ability to avoid the predators developed in later generations, then selection on wing size and shape would potentially become far weaker. Study systems like the one used here allow for additional future work to address these questions in a relatively straightforward manner, which will likely become increasingly important as we recognize the limitations of measuring relatively small numbers of traits.

**Table 1.** Survival ability in the base population measured as the number of surviving flies in each arena after 24 hours exposure with the predators. Table shows the output from the lm funtion in R for a set of models evaluated using Bayesian information criteria. Numbers in parenthesis are standard errors of the above estimates.

|  | Model 1 | Model 2 | Model 3 |
| --- | --- | --- | --- |
| (Intercept) | 6.71<br>(0.38) | 7.14<br>(0.50) | 7.33<br>(0.60) |
| Sex: M | -2.91***<br>(0.55) | -2.94***<br>(0.55) | -3.33***<br>(0.84) |
| Date: June 2008 |  | -0.77<br>(0.59) | -1.03<br>(0.82) |
| Date: March 2008 |  | -0.57<br>(0.89) | -1.33<br>(1.40) |
| Sex: M x Date: June 2008 |  |  | 0.53<br>(1.20) |
| Sex: M x Date: March 2008 |  |  | 1.33<br>(1.84) |
| R<sup>2</sup> | 0.42 | 0.45 | 0.46 |
| Adj. R<sup>2</sup> | 0.41 | 0.40 | 0.38 |
| Num. obs. | 41 | 41 | 41 |
| ΔAIC | 0.0 | 3.1 | 8.1 |
| ΔBIC | 0.0 | 5.5 | 12.2 |
| ΔDeviance | 7.4 | 1.8 | 0.0 |
| BIC Weights | 0.937 | 0.061 | 0.002 |

\*\*\* $p < 0.01$ , \*\* $p < 0.05$ , \* $p < 0.1$

**Table 2.**
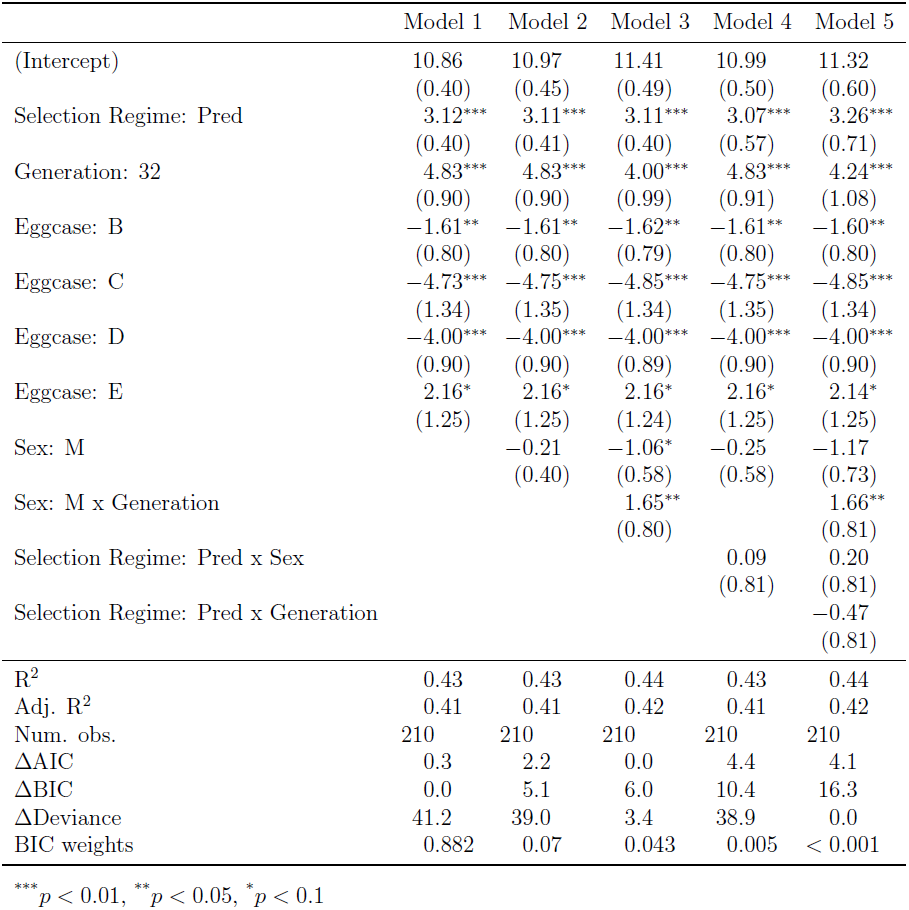
Survival ability in the evolved populations measured as the number of surviving flies in each arena after 24 hours exposure with the predators. Table shows the output from the lm funtion in R for a set of models evaluated using Bayesian information criteria. Numbers in parenthesis are standard errors of the above estimates.

|  | Model 1 | Model 2 | Model 3 | Model 4 | Model 5 |
| --- | --- | --- | --- | --- | --- |
| (Intercept) | 10.86<br>(0.40) | 10.97<br>(0.45) | 11.41<br>(0.49) | 10.99<br>(0.50) | 11.32<br>(0.60) |
| Selection Regime: Pred | 3.12***<br>(0.40) | 3.11***<br>(0.41) | 3.11***<br>(0.40) | 3.07***<br>(0.57) | 3.26***<br>(0.71) |
| Generation: 32 | 4.83***<br>(0.90) | 4.83***<br>(0.90) | 4.00***<br>(0.99) | 4.83***<br>(0.91) | 4.24***<br>(1.08) |
| Eggcase: B | -1.61**<br>(0.80) | -1.61**<br>(0.80) | -1.62**<br>(0.79) | -1.61**<br>(0.80) | -1.60**<br>(0.80) |
| Eggcase: C | -4.73***<br>(1.34) | -4.75***<br>(1.35) | -4.85***<br>(1.34) | -4.75***<br>(1.35) | -4.85***<br>(1.34) |
| Eggcase: D | -4.00***<br>(0.90) | -4.00***<br>(0.90) | -4.00***<br>(0.89) | -4.00***<br>(0.90) | -4.00***<br>(0.90) |
| Eggcase: E | 2.16*<br>(1.25) | 2.16*<br>(1.25) | 2.16*<br>(1.24) | 2.16*<br>(1.25) | 2.14*<br>(1.25) |
| Sex: M |  | -0.21<br>(0.40) | -1.06*<br>(0.58) | -0.25<br>(0.58) | -1.17<br>(0.73) |
| Sex: M x Generation |  |  | 1.65**<br>(0.80) |  | 1.66**<br>(0.81) |
| Selection Regime: Pred x Sex |  |  |  | 0.09<br>(0.81) | 0.20<br>(0.81) |
| Selection Regime: Pred x Generation |  |  |  |  | -0.47<br>(0.81) |
| R<sup>2</sup> | 0.43 | 0.43 | 0.44 | 0.43 | 0.44 |
| Adj. R<sup>2</sup> | 0.41 | 0.41 | 0.42 | 0.41 | 0.42 |
| Num. obs. | 210 | 210 | 210 | 210 | 210 |
| ΔAIC | 0.3 | 2.2 | 0.0 | 4.4 | 4.1 |
| ΔBIC | 0.0 | 5.1 | 6.0 | 10.4 | 16.3 |
| ΔDeviance | 41.2 | 39.0 | 3.4 | 38.9 | 0.0 |
| BIC weights | 0.882 | 0.07 | 0.043 | 0.005 | < 0.001 |

\*\*\* $p < 0.01$ , \*\* $p < 0.05$ , \* $p < 0.1$

**Table 3.**
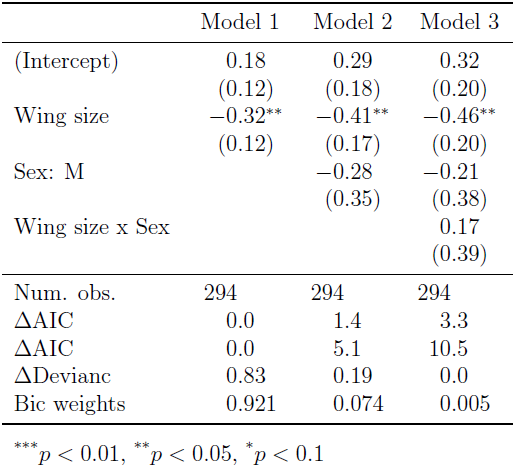
The output from the logistic regression of wing size onto survival in the base population for a set of models evaluated using Bayesian information criteria. Numbers in parenthesis are standard errors of the above estimates. The logistic regression models were used to evaluate statistical significance of the estimated selection differentials. The values reported in the manuscript were taken from identical linear regression models (output not shown)

**Table 4.** The output from the logistic regression of wing size onto survival in the evolved populations for a set of models evaluated using Bayesian information criteria. Numbers in parenthesis are standard errors of the above estimates.The logistic regression models were used to evaluate statistical significance of the estimated selection differentials. The values reported in the manuscript were taken from identical linear regression models (output not shown)

|  | Model 1 | Model 2 | Model 3 | Model 4 | Model 5 |
| --- | --- | --- | --- | --- | --- |
| (Intercept) | 0.50<br>(0.11) | 0.53<br>(0.11) | 0.64<br>(0.12) | 0.86<br>(0.15) | 0.79<br>(0.16) |
| Wing size | -0.42***<br>(0.09) | -0.35***<br>(0.09) | -0.40***<br>(0.09) | -0.62***<br>(0.13) | -0.55***<br>(0.14) |
| Selection Regime: Pred | 0.55***<br>(0.07) | 0.31***<br>(0.09) | 0.16<br>(0.11) | 0.16<br>(0.11) | 0.16<br>(0.11) |
| Replicate: R2 | -0.27***<br>(0.10) | -0.60***<br>(0.13) | -0.57***<br>(0.13) | -0.57***<br>(0.13) | -0.57***<br>(0.13) |
| Sex: M | -0.45***<br>(0.17) | -0.29*<br>(0.18) | -0.34*<br>(0.18) | -0.73***<br>(0.23) | -0.76***<br>(0.23) |
| Generation: G32 | -0.48***<br>(0.11) | -0.43***<br>(0.11) | -0.63***<br>(0.14) | -1.02***<br>(0.20) | -1.01***<br>(0.20) |
| Wing size x Selection Regime | 0.19***<br>(0.07) | 0.20***<br>(0.07) | 0.22***<br>(0.07) | 0.24***<br>(0.07) | 0.23***<br>(0.07) |
| Replicate: R2 x Generation | 0.96***<br>(0.14) | 1.02***<br>(0.14) | 1.03***<br>(0.14) | 1.09***<br>(0.15) | 1.09***<br>(0.15) |
| Sex M x Generation | 0.69***<br>(0.14) | 0.64***<br>(0.14) | 0.64***<br>(0.14) | 1.42***<br>(0.33) | 1.38***<br>(0.33) |
| Selection Regime: Pred x Replicate |  | 0.60***<br>(0.15) | 0.58***<br>(0.15) | 0.53***<br>(0.15) | 0.52***<br>(0.15) |
| Selection Regime: Pred x Generation |  |  | 0.35**<br>(0.14) | 0.32**<br>(0.14) | 0.33**<br>(0.14) |
| Wing size x Generation |  |  |  | 0.44***<br>(0.17) | 0.44***<br>(0.17) |
| Wing size x Sex |  |  |  |  | -0.19<br>(0.17) |
| Num. obs. | 3932 | 3932 | 3932 | 3932 | 3932 |
| ΔAIC | 23.6 | 8.9 | 4.8 | 0.0 | 0.7 |
| ΔBIC | 8.4 | 0.0 | 2.2 | 3.6 | 10.6 |
| ΔDeviance | 30.95 | 14.25 | 8.12 | 1.3 | 0.0 |
| BIC weights | 0.001 | 0.658 | 0.224 | 0.108 | > 0.001 |

\*\*\* $p < 0.01$ , \*\* $p < 0.05$ , \* $p < 0.1$

**Table 5.**
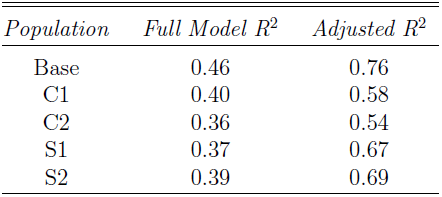
R^2^ values from partialleast squares analysis for shape.

| *Population* | *Full Model $R^2$* | *Adjusted $R^2$* |
| --- | --- | --- |
| Base | 0.46 | 0.76 |
| C1 | 0.40 | 0.58 |
| C2 | 0.36 | 0.54 |
| S1 | 0.37 | 0.67 |
| S2 | 0.39 | 0.69 |

## Acknowledgments

This manuscript was improved by comments from Jeff Conner, Kay Holekamp, Barry Williams, Tony Frankino, and members of the Dworkin lab. This material is based in part upon work supported by NIH grant 1R01GM09442401 (to ID). This material is based in part upon work supported by the National Science Foundation under Cooperative Agreement No. DBI-0939454. Any opinions, findings, and conclusions or recommendations expressed in this material are those of the author(s) and do not necessarily reflect the views of the National Science Foundation.

---

**Figure S1.**
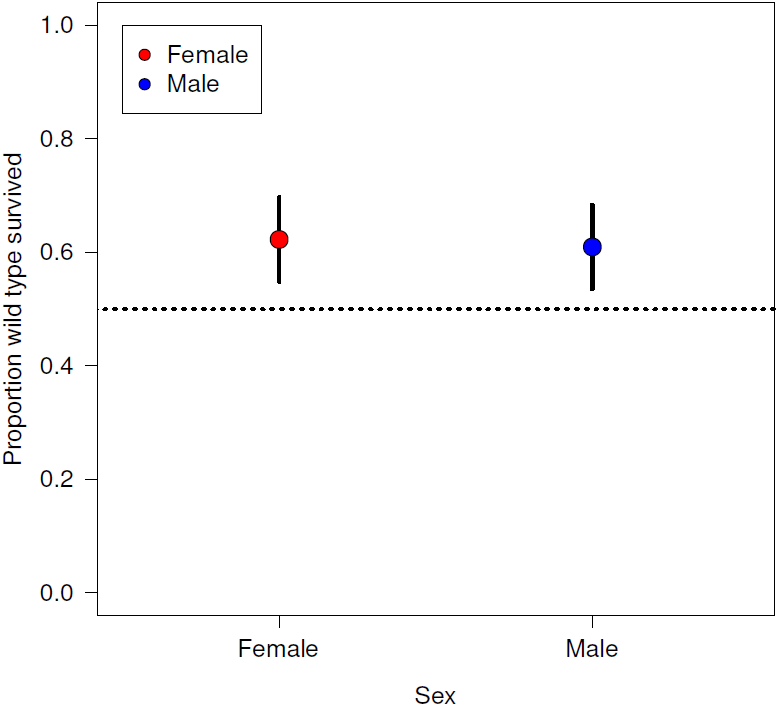
Proportion of wild-type flies surviving in each arena. Error bars are 95% confidence intervals.

**Figure S2.**
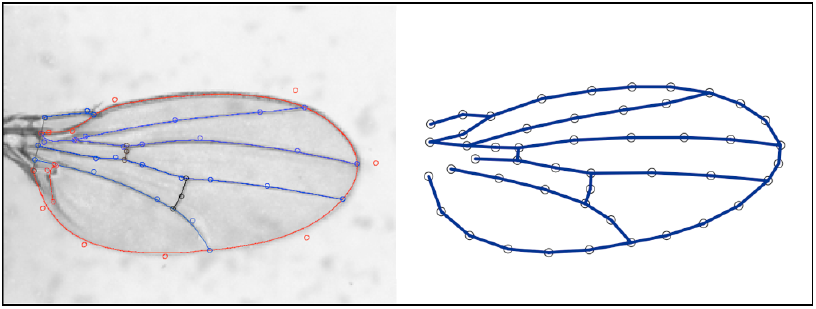
A greyscale wing image from a microscope-mounted camera, showing the splines fitted by the WingMachine software (left panel). Landmarks and semi-landmarks can be extracted from these splines, and superimposed using the CPReader program. From this data the wing is modelled by the position of these landmarks and semi-landmarks (represented by the open circles in the right panel), and simply connecting these coordinates with straight line segments as in the right panel allows for an easily interpretable visualisation of the wing veins and margins.

**Figure S3.**
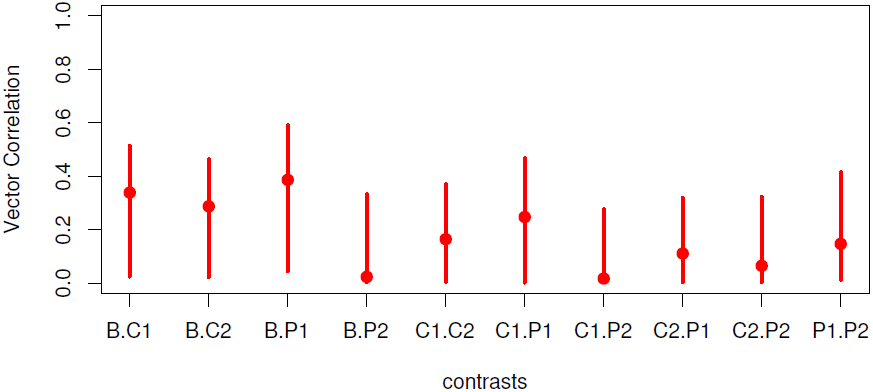
Vector correlations between selection differentials (**S**) measured in the base population (B) and all four experimental evolution populations. Points and lines are vector correlations and bootstrapped 95% confidence intervals.

**Figure S4.**
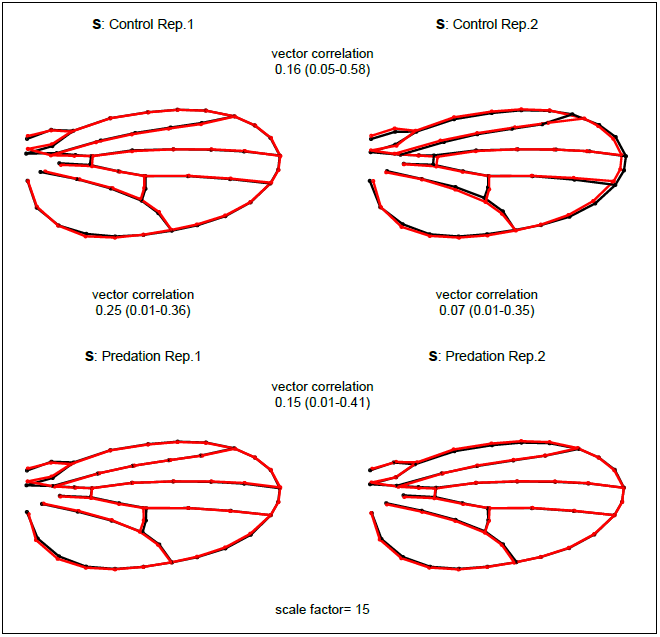
Visualisation of the selection differential (S) as measured at generations 31 & 32 (see methods) in the experimental evolution populations. The shapes represent the mean shape plus15x S (black line) and minus 15x **S** (red line), and the points represent landmarks and semi-landmarks. Vector correlations between **S** vectors (and their 95% credible intervals) are printed between the pairs of populations to which they relate.

